# Sensitivity analysis based ranking reveals unknown biological hypotheses for down regulated genes in time buffer during administration of PORCN-WNT inhibitor ETC-1922159 in CRC^†^

**DOI:** 10.1101/180927

**Authors:** Shriprakash Sinha

## Abstract

In a recent development of the PORCN-WNT inhibitor ETC-1922159 for colorectal cancer, a list of down-regulated genes were recorded in a time buffer after the administration of the drug. The regulation of the genes were recorded individually but it is still not known which higher (≥ 2) order interactions might be playing a greater role after the administration of the drug. In order to reveal the priority of these higher order interactions among the down-regulated genes or the likely unknown biological hypotheses, a search engine was developed based on the sensitivity indices of the higher order interactions that were ranked using a support vector ranking algorithm and sorted. For example, LGR family (Wnt signal enhancer) is known to neutralize RNF43 (Wnt inhibitor). After the administration of ETC-1922159 it was found that using HSIC (and rbf, linear and laplace variants of kernel) the rankings of the interaction between LGR5-RNF43 were 61, 114 and 85 respectively. Rankings for LGR6-RNF43 were 1652, 939 and 805 respectively. The down-regulation of LGR family after the drug treatment is evident in these rankings as it takes bottom priorities for LGR5-RNF43 interaction. The LGR6-RNF43 takes higher ranking than LGR5-RNF43, indicating that it might not be playing a greater role as LGR5 during the Wnt enhancing signals. These rankings confirm the efficacy of the proposed search engine design. Conclusion: Prioritized unknown biological hypothesis form the basis of further wet lab tests with the aim to reduce the cost of (1) wet lab experiments (2) combinatorial search and (3) lower the testing time for biologist who search for influential interactions in a vast combinatorial search forest. From in silico perspective, a framework for a search engine now exists which can generate rankings for *n*^*th*^ order interactions in Wnt signaling pathway, thus revealing unknown/untested/unexplored biological hypotheses and aiding in understanding the mechanism of the pathway. The generic nature of the design can be applied to any signaling pathway or phenomena under investigation where a prioritized order of interactions among the involved factors need to be investigated for deeper understanding. Future improvements of the design are bound to facilitate medical specialists/oncologists in their respective investigations.

**Significance:** Recent development of PORCN-WNT inhibitor enantiomer ETC-1922159 cancer drug show promise in suppressing some types of colorectal cancer. However, the search and wet lab testing of unknown/unexplored/untested biological hypotheses in the form of combinations of various intra/ extracellular factors/genes/proteins affected by ETC-1922159 is not known. Currently, a major problem in biology is to cherry pick the combinations based on expert advice, literature survey or guesses to investigate a particular combinatorial hypothesis. A search engine has be developed to reveal and prioritise these unknown/untested/unexplored combinations affected by the inhibitor. These ranked unknown biological hypotheses facilitate in narrowing down the investigation in a vast combinatorial search forest of ETC-1922159 affected synergistic-factors.

## Introduction

### Wnt signaling and secretion

Sharma^1^’s accidental discovery of the Wingless played a pioneering role in the emergence of a widely expanding research field of the Wnt signaling pathway. A majority of the work has focused on issues related to • the discovery of genetic and epigenetic factors affecting the pathway Thorstensen *et al*.^2^ & Baron and Kneissel^3^, • implications of mutations in the pathway and its dominant role on cancer and other diseases Clevers^4^, • investigation into the pathway’s contribution towards embryo development Sokol^5^, homeostasis Pinto *et al*^6^ & Zhong *et al*.^7^ and apoptosis Pećina-Šlaus^8^ and • safety and feasibility of drug design for the Wnt pathway Kahn^9^, Garber^10^, Voronkov and Krauss^11^, Blagodatski *et al*.^12^ & Curtin and Lorenzi^13^.

The Wnt phenomena can be roughly segregated into signaling and secretion part. The Wnt signaling pathway works when the WNT ligand gets attached to the Frizzled(FZD)/LRP coreceptor complex. FZD may interact with the Dishevelled (DVL) causing phosphorylation. It is also thought that Wnts cause phosphorylation of the LRP via casein kinase 1 (CK1) and kinase GSK3. These developments further lead to attraction of Axin which causes inhibition of the formation of the degradation complex. The degradation complex constitutes of AXIN, the *β*-catenin transportation complex APC, CK1 and GSK3. When the pathway is active the dissolution of the degradation complex leads to stabilization in the concentration of *β*-catenin in the cytoplasm. As *β*-catenin enters into the nucleus it displaces the GROUCHO and binds with transcription cell factor TCF thus instigating transcription of Wnt target genes. GROUCHO acts as lock on TCF and prevents the transcription of target genes which may induce cancer. In cases when the Wnt ligands are not captured by the coreceptor at the cell membrane, AXIN helps in formation of the degradation complex. The degradation complex phosphorylates *β*-catenin which is then recognised by F BOX/WD repeat protein *β*-TRCP. *β*-TRCP is a component of ubiquitin ligase complex that helps in ubiquitination of *β*-catenin thus marking it for degradation via the proteasome. A cartoon of the signaling transduction snapshot is shown in figure 1.

**Fig. 1.**
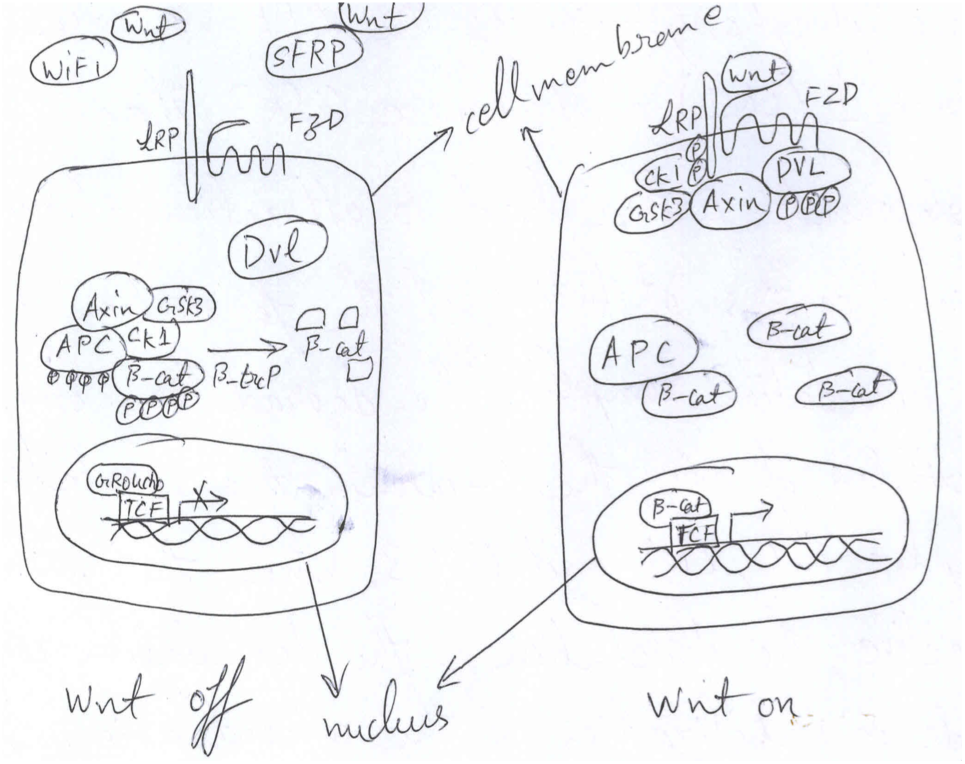
Cartoon of Wnt Signaling.

Contrary to the signaling phenomena, the secretion phenomena is about the release and transportation of the WNT protein/ligand in and out of the cell, respectively. Briefly, the WNT proteins that are synthesized with the endoplasmic reticulum (ER), are known to be palmitoyleated via the Porcupine (PORCN) to form the WNT ligand, which is then ready for transportation Tanaka *et al*.^14^. It is believed that these ligands are then transported via the EVI/WNTLESS transmembrane complex out of the cell Bänziger *et al*.^15^ & Bartscherer *et al*.^16^. The EVI/WNTLESS themselves are known to reside in the Golgi bodies and interaction with the WNT ligands for the later’s glycosylation Kurayoshi *et al*.^17^ & Gao and Hannoush^18^. Once outside the cell, the WNTs then interact with the cell receptors, as explained in the foregoing paragraph, to induce the Wnt signaling. Of importance is the fact that the EVI/WNTLESS also need a transporter in the from of a complex termed as Retromer. A cartoon of the signaling transduction snapshot is shown in figure 2.

**Fig. 2.**
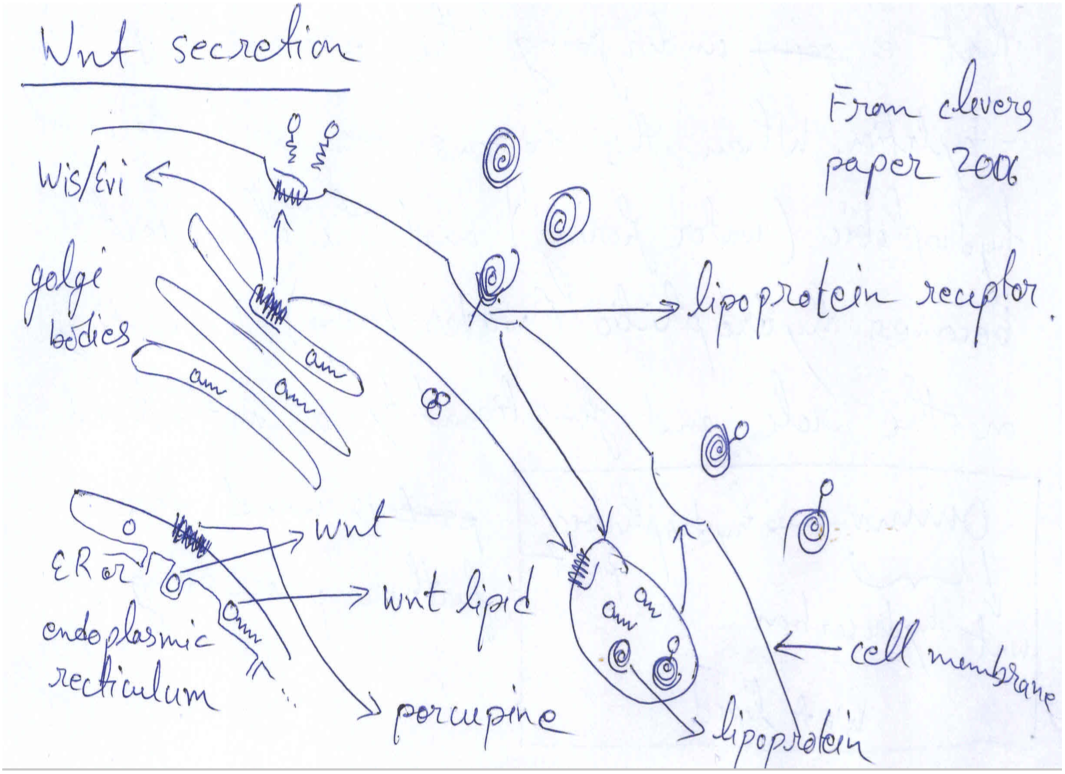
Cartoon of Wnt Secretion.

### PORCN-WNT inhibitors

The regulation of the Wnt pathway is dependent on the production and secretion of the WNT proteins. Thus, the inhibition of a causal factor like PORCN which contributes to the WNT secretion has been proposed to be a way to interfere with the Wnt cascade, which might result in the growth of tumor. Several groups have been engaged in such studies and known PORCN-WNT inhibitors that have been made available till now are IWP-L6 Chen *et al*.^19^ & Wang *et al*. ^20^, C59 Proffitt *et al*. ^21^, LGK974 Liu *et al*. ^22^ and ETC-1922159 Duraiswamy *et al*. ^23^. In this study, the focus of the attention is on the implications of the ETC-1922159, after the drug has been administered. The drug is a enantiomer with a nanomolar activity and excellent bioavailability as claimed in Duraiswamy *et al*. ^23^.

## Methodology

Note that the technical details employed to derive results have already been described in a parallel manuscript submitted to this journal. Interested readers are advised to go through the manuscript titled "Prioritizing 2^*nd*^ & 3 ^*rd*^ order interactions via support vector ranking using sensitivity indices on static/time series Wnt measurements" by this author Sin^24^.

### Revealing higher order biological hypotheses via sensitivity analysis and insilico ranking algorithm

In the trial experiments on ETC-1922159 Madan *et al*. ^25^, a list of genes (2500±) have been reported to be up and down regulated after the drug treatment and a time buffer of 3 days. Some of the transcript levels of these genes have been recorded and the experimental design is explained elaborately in the same manuscript. In the list are also available unknown or uncharacterised proteins that have been recorded after the drug was administered. These have been marked as "‐ ‐" in the list (Note - In this manuscript these uncharacterised proteins have been marked as "XXM", were *M* = 1,2,3,…). The aim of this work is to reveal unknown/unexplored/untested biological hypotheses that form higher order combinations. For example, it is known that the combinations of WNT-FZD or RSPO-LGR-RNF play significant roles in the Wnt pathway. But the *n* ≥ 2,3,…-order combinations out of *N*(> *n*) genes forms a vast combinatorial search forest that is extremely tough to investigate due to the humongous amount of combinations. Currently, a major problem in biology is to cherry pick the combinations based on expert advice, literature survey or random choices to investigate a particular combinatorial hypothesis. The current work aims to reveal these unknown/unexplored/untested combinations by prioritising these combinations using a potent support vector ranking algorithm Joachims^26^. This cuts down the cost in time/energy/investment for any investigation concerning a biological hypothesis in a vast search space.

The pipleline works by computing sensitivity indicies for each of these combinations and then vectorising these indices to connote and form discriminative feature vector for each combination. The ranking algorithm is then applied to a set of combinations/sensitivity index vectors and a ranking score is generated. Sorting these scores leads to prioritization of the combinations. Note that these combinations are now ranked and give the biologists a chance to narrow down their focus on crucial biological hypotheses in the form of combinations which the biologists might want to test. Analogous to the webpage search engine, where the click of a button for a few key-words leads to a ranked list of web links, the pipeline uses sensitivity indices as an indicator of the strength of the influence of factors or their combinations, as a criteria to rank the combinations. The generic pipleline for the generation for ranking has been shown in figure 3 from one of the author’s unpublished/submitted manuscript Sin^24^, the time series data set for which was obtained from Gujral and MacBeath^27^.

**Fig. 3.**
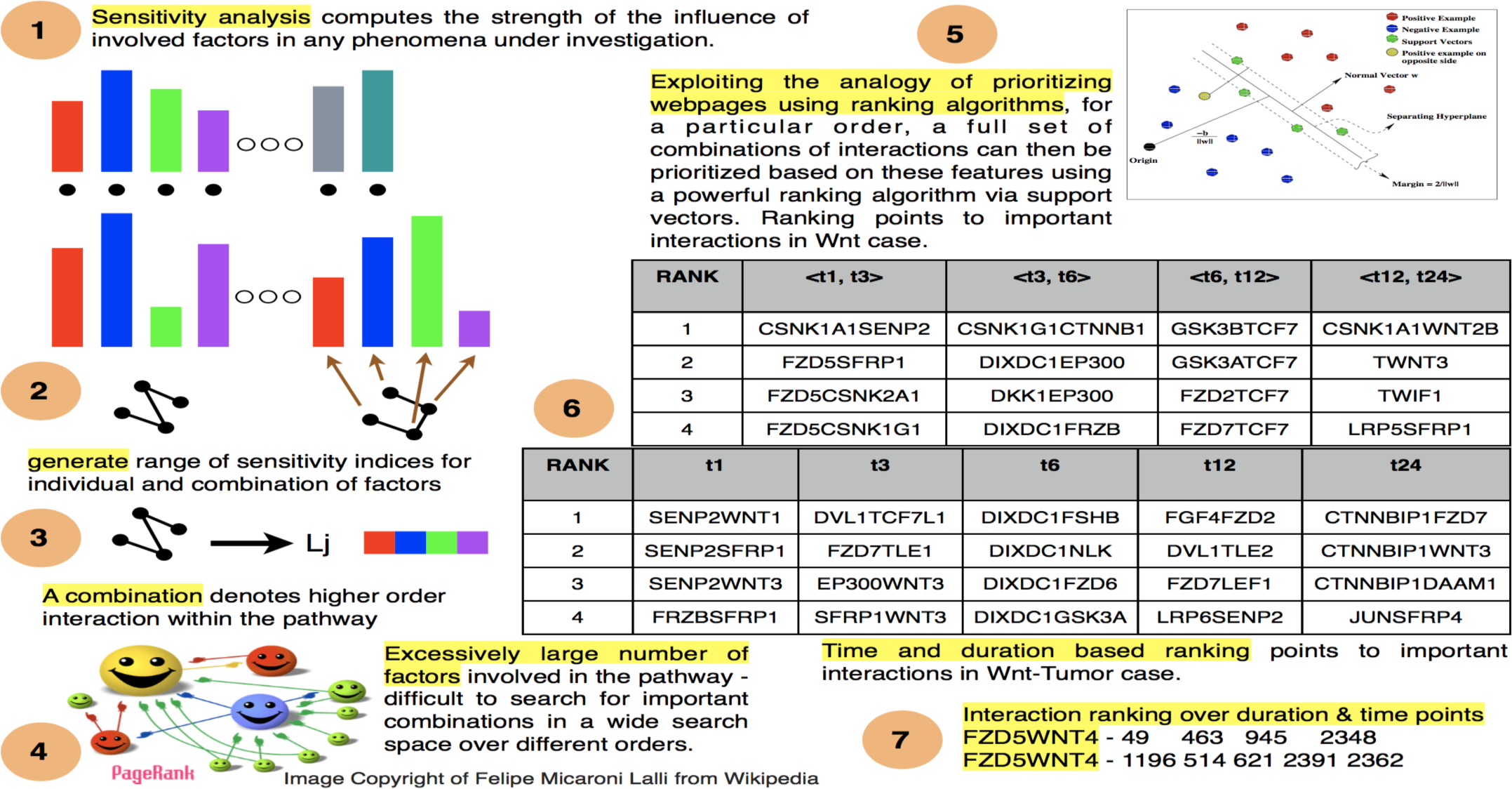
Graphical abstract of how the whole search engine works to find out crucial interactions within the components of the pathway under
consideration as different time points and durations. Figure adopted from a manuscript titled "Prioritizing 2^*nd*^& 3^*rd*^ order interactions via support vector ranking using sensitivity indices on static/time series Wnt measurements", that has been submitted in parallel by this author Sin ^24^.

### Translating the pipeline into code of execution in R

The pipeline depicted in figure 3 was modified and translated into code in R programming language R Development Core Team^28^. Figure 4 shows one such computation and the main commands and some of the internal executions are shown in red. The execution begins by some preprocessing of the file containing the information regarding the down regulated genes via *source*(”*extractETCdata.R*”). The library containing the sensitivity analysis methods is then loaded in the interface using the command *library*(*sensitivity*) Faivre *et al*. ^29^ & Iooss and Lemaître^30^. Next a gaussian distribution for a particular transcript level for a gene is generated to have a sample. This is needed in the computation of sensitivity index for that particular gene. A gene of particular choice is selected (in blue) and the choice of the combination is made (here k = 2). Later a kernel is used (here rbf for HSIC method). 50 different indices are generated which help in generating aggregate rank using these 50 indices. The Support vector ranking algorithm is employed to generate the scores for different combinations and these scores are then sorted out. This is done using the command *source* (”*SV MRank* – *Results* – *S* – *mean.R*”). A file is generated that contains the combinations in increasing order of influence after the ETC-1922159 has been administered. Note that 1 means lowest rank and 2743 is the highest rank for 2^*nd*^ order combinations for gene RAD51AP1 using HSIC-rbf density based sensitivity index. The code has been depicted in figure 4.

**Fig. 4.**
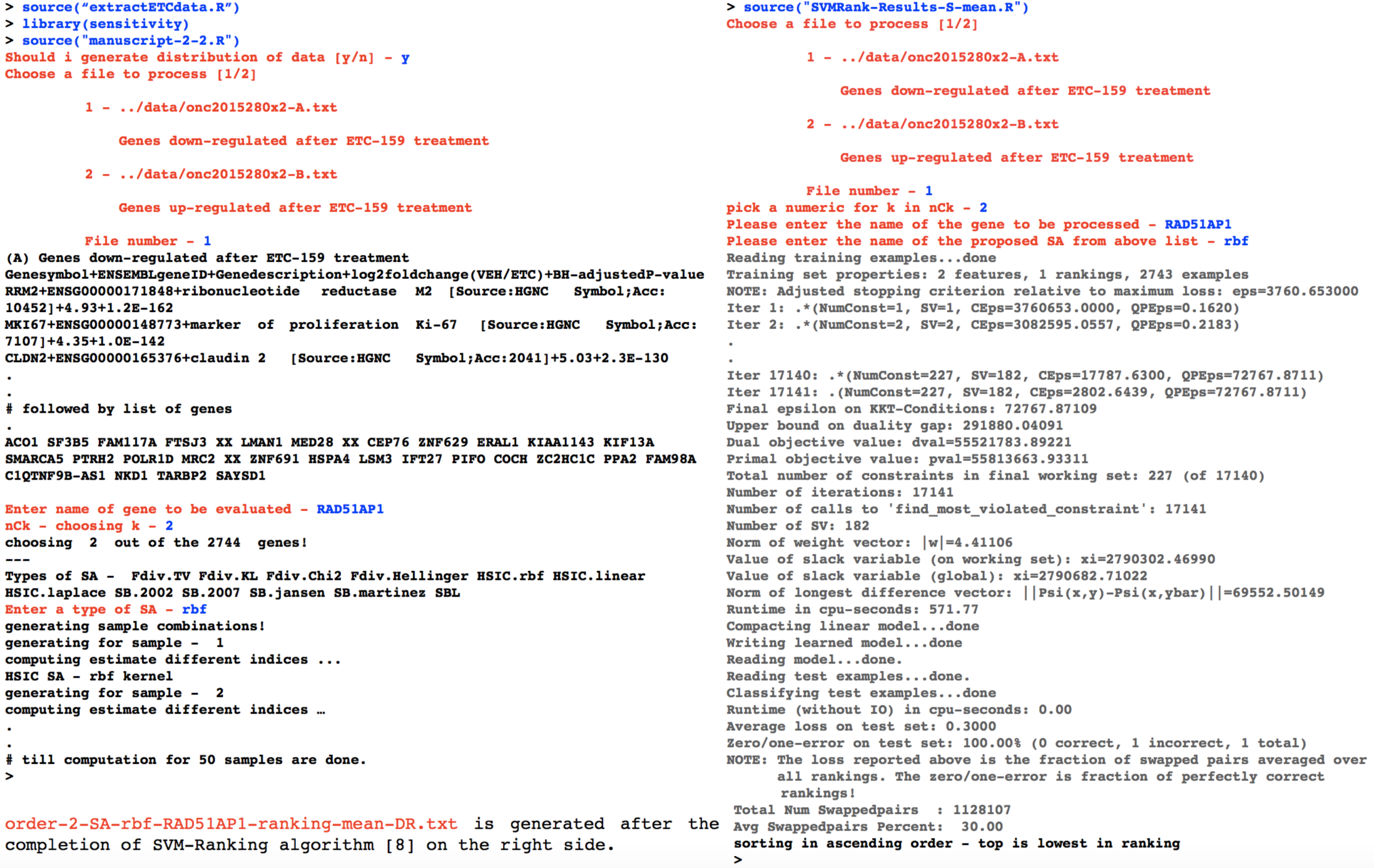
Translation of a modification of the pipeline in figure 3 into code of execution in R.

### Interpretation of 2^*nd*^ order ranking with RAD51AP1 using HSIC-rbf density based sensitivity index

For example, the ranking generated for second order combination for RAD51AP1 is enlisted below. What these rankings suggests is that after the treatment of ETC-1922159, genes along with their combination with RAD51AP1 were prioritised and this gives the biologists a clue to study the behaviour of RAD51AP1 along with other genes in colorectal cancer case.

1 HOXB9-RAD51AP1
2 NASP-RAD51AP1
3 MCM3-RAD51AP1

•

•

2741 RAD51AP1-CDX2

2742 RAD51AP1-LPCAT3

2743 RAD51AP1-EIF2B1

DNA repair is an important aspect in maintaining the proper and healthy functioning for the cells in the human body. Failure in DNA repair process can lead to aberrations as well as tumorous stages. There are various types of damages that a DNA can go through, one of which is the DNA double strand breaks (DSB) that can be repaired via homologous recombination (HR). RAD51 plays a central role in HR and is known to function in the three phases of HR namely : presynapsis, synapsis and post-synapsis Krejci *et al*.^31^. Recently, RAD51 has been implicated as a negative/poor prognostic marker for colorectal adenocarcinoma and has been found to be highly expressed Tennstedt *et al*. ^32^. A negative/poor prognostic marker indicates that it is harder to control the malignancy. Since RAD51 helps in the repair of the DNA damage via HR and is implicated as a poor prognostic marker in colorectal adenocarcinoma, this suggests its functionality in maintaining genomic stability and therapeutic resistance to cancer drugs. Mechanistically, RAD51AP1 facilitates RAD51 during the repairing process by binding with RAD51 via two DNA binding sites, thus helping in the D-loop formation in the HR process Dunlop *et al*.^33^ & Modesti *et al*.^34^.

In context of the ETC-1922159 treatment, a series of 2^*nd*^ order rankings have been generated for RAD51AP1. Using the above pipeline can help decipher biological implications. BRCA2 is known to be a main mediator in the HR process along with RAD51 and interact with RAD51 in various ways Davies and Pellegrini^35^, Esashi *et al*.^36^, Ayoub *et al*.^37^ & Schlacher *et al*.^38^. A relative low rank of 458 between RAD51AP1-BRCA2 states that after the ETC-1922159 treatment the combination gets low priority indicating that as there is suppression of CRC, the genomic stability of the cancer cells is affected by rendering the DNA repairing capacity of RAD51AP1-BRCA2 ineffective to a certain extent. PPA2 is a tumor suppressor that regulates many signaling pathways and plays crucial role in cell transformation Seshacharyulu *et al*. ^39^. PPA2 in known to be highly suppressed in colorectal cancer case Cristóbal *et al*.^40^. The relatively high ranking of 2675 for RAD51AP1-PPA2 combination indicates two points (a) the ineffectiveness of RAD51AP1 to provide genomic stability to cancer cell and (b) the over expression of PPA2 after ETC-1922159 was administered. Also, SET is a known PPA2 inhibitor and is observed to be highly overexpressed in colorectal cancer cases. In relation to the ranking of RAD51AP1-PPA2, after the administration of the drug, RAD51AP1-SET showed a lower rank of 2227.

The x-ray repair cross complementing XRCC family is known to work as a mediator or stabilizer for RAD51 during the HR process Krejci *et al*.^31^. The relatively low rank of 366 for the RAD51AP1-XRCC2 indicates that the effectiveness for DNA repair capability is affected by ETC-1922159 drug as the tumor growth is suppressed. This is further confirmed by the fact that XRCC2 forms complex with paralogues of RAD51, i.e RAD51C—RAD51D—XRCC2 for DNA repair Masson *et al*.^41^ & Yokoyama *et al*.^42^. Other members of XRCC like XRCC1, XRCC6BP1 and XRCC6, showed relatively higher rankings of 1874, 2398 and 2558. Not much is known about these 3 factors in colorectal cancer case. XRCC1 may be weakly implicated in the colorectal cancer cases as have been found in the case studies in Taiwan Yeh *et al*.^43^. Very little is known about XRCC6BP1 and in a risk score based analysis it was found that XRCC6BP1 acts as a tumor repressor with very low expression profile in colorectal cancer Diao *et al*.^44^. Hight rankings suggests that after ETC-1922159 treatment, the expression level of XRCC6BP1 is extremely high, indicating the establishment of genomic stability of health via suppression of cancer cells. Finally, the very high priority of XRCC6 only indicates its role as a tumor repressor and further wet lab analysis needs to be conducted to verify the effectiveness of the pipeline.

RNF43/ZNRF3 are known to negatively regulate the Wnt pathway by targeting the FZD family and leading to its degradation and thus acting as a hinderance to the Wnt pathway Feng and Gao^45^. In presence of members from RSPO and LGR family, the RNF43/ZNRF3 is degraded in the cell and this leads to enhancement of the Wnt signaling de Lau *et al*. ^46^. In relation to the RAD51AP1, the ranking of RNF43 was found to be 1893 and the ranking of ZNRF3 was found to be 549, respectively, after the treatment of ETC-1922159. The lower ranking of ZNRF43 with RAD51AP1 might indicate some affinity between RAD51AP1-ZNRF43 in comparison to the high ranking of RAD51AP1-RNF43. This needs to be tested. LGR5 and LGR6 had associated low ranks for 327 and 161, respectively. After the administration of the drug, the LGR5, RNF43 and ZNRF43 were known to be downregulated Madan *et al*. ^25^. These are evident from the low rankings in combination with RAD51AP1, except for RNF43. An implication of the above combination of rankings can be the fact that as the drug takes its affect, it has multiple consequences wherein the RNF43/ZNRF3 along with LGR5 is being degraded so that the signaling is inhibited and also the genomic instability of the cancer cells in instigated by the ineffectiveness of RAD51AP1 to help in DNA repair in cancer cells.

Such rankings hold promise for any biologists who is investigating a pathway for a particular phenomena and is faced with a vast combinatorial search forest. These rankings also provide confirmatory results for existing published wet lab affirmations.

## Results & discussion

### MYC-HOXB8-EZH2

EZH2 encodes enhancer of zeste homolog 2 and is involved in transcriptional repression via epigenetic modifications. It has been found to be either mutated or over-expressed in many forms of cancer. Over expression of EZH2 leads to silencing of various tumor suppressor genes and thus implicating it for potential roles in tumorigenesis^47^. EZH2 is a subunit of the highly conserved Polycomb repressive complex 2 (PRC2) which executes the methylation of the histone H3 at lysine-27^48^. Thus targeting EZH2 has become a major research domain for cancer therapeutics ^49^. In colon cancer, it has been shown that depletion of EZH2 has led to blocking of proliferation of the cancer^50^. This indicates the fact that tumor suppressor genes get activated and lead to subsequent blocking of the cancer. Also, EZH2 is recruited by PAF to bind with *β*-catenin transcriptional complex for further Wnt target gene activation, independent of the EZH2 epigenetic modification activities^51^.

Consistent with these, ETC-15922159 treatment lead to down regulation of EZH2 in colorectal cancer samples^25^. This would have activated a lot of tumor suppressor genes that led to subsequent suppression of regrowth in treated cancer samples. More importantly, MYC directly upregulates core components of PRC2, EZH2 being one of them, in embryonic stem cells^52^.^52^ show that silencing of c-MYC and N-MYC ^53^ lead to reduction in the expression of PRC2 and thus EZH2. Furthermore, in colorectal cancer cases, ^54^ show that knockdown of MYC led to decrease in EZH2 levels. Similar findings have been observed in^55^,^56^ &^57^. Our in silico findings show consistent results with respect to this down regulation after assigning a low rank of 54 along with MYC-HOXB8.

More specifically, our in silico pipeline is able to approximate the value of the 3^rd^ order combination of MYC-HOXB8-EZH2 by assigning a rank that is consistent with wet lab findings of dual combinatorial behaviour of MYC-EZH2 and MYC-HOXB8. However, the since the mechanism of combination of MYC-HOXB8 is not known hitherto, it would be interesting to confirm the behaviour of MYC-HOXB8-EZH2 at 3^rd^ order to reveal a portion of the Wnt pathway’s modus operandi in colorectal cancer. Further wet lab tests on these in silico findings will confirm the efficacy of the search engine.

### Identified/identified 2^*nd*^-order combinations

Table 1 shows the relative rankings apropos RNF43 for three different kernels. The genes that have been shown along with the RNF43 are known to be highly expressed in colorectal cancer cases and after the ETC-1922159 was administered. Here we show only a few of the rankings as a confirmatory result that supports the in silico findings of the pipeline. In the table, except for BMP7 and NKD1, while considering the majority rankings over the three different columns representing the kinds of kernels used with the HSIC density based method, we find that each one of them has been assigned a relatively low priority in the order of 2743 combinations. The implication of these low rankings indicate or confirm the effectiveness of the pipeline in assigning appropriate biologically induced priority to the ETC-1922159 influenced down regulation of these genes. Surprisingly, BMP7 Motoyama *et al*. ^58^ and NKD1 Stancikova *et al*. ^59^ are known to be highly expressed in colorectal cancer case and were down regulated after the drug treatment. But these were assigned high rank with RNF43. Mutations in RNF43 could be subdued after ETC-1922159 has suppressed the Wnt pathway as has been shown in Madan *et al*. ^25^. NKD1 is enigmatic in nature and known to be a negative regulator of the Wnt pathway Larraguibel *et al*. ^60^. Mutations in NKD1 have been found to be prevalent in colorectal cancer cases Guo *et al*. ^61^. High rank might suggest that after the suppression of the cancer cells, the NKD1 is highly activated or the mutated version of NKD1, if present, are highly ineffective. Thus the RNF43-NKD1 acquires a high ranking by the engine and it is more likely that NKD1 is highly activated.

**Table 1.** 2^*nd*^ order interaction ranking using HSIC for different kernels. Total number of interactions per gene were 2743.

| RANKING USING HSIC W.R.T RNF43 |  |  |  |
| --- | --- | --- | --- |
| 2<sup>nd</sup> odr comb. | rbf | laplace | linear |
| LGR5-RNF43 | 61 | 85 | 114 |
| LGR6-RNF43 | 1652 | 805 | 939 |
| RAD51AP1-RNF43 | 175 | 469 | 11 |
| ZNRF3-RNF43 | 570 | 1603 | 404 |
| RRM2-RNF43 | 556 | 667 | 442 |
| MKI67-RNF43 | 1575 | 278 | 92 |
| AXIN2-RNF43 | 531 | 51 | 482 |
| ASCL2-RNF43 | 116 | 441 | 197 |
| MCM4-RNF43 | 131 | 1142 | 204 |
| RNF43-BMP7 | 2723 | 2701 | 2530 |
| RNF43-NKD1 | 2635 | 2169 | 2226 |

ZNRF3/RNF43 is known to be implicated in inhibiting the Wnt signaling via degradation of FZDs at the cell surface level de Lau *et al*. ^46^ & Hao *et al*. ^62^. Mutations in these can lead to abberent signaling and mutated RNF43 were found to be suppressed after the ETC-1922159 treatment and the relatively low rank assigned to the combination points to effectiveness of the pipeline. Similarly, RRM2 which is involved in DNA repair Zhang *et al*. ^63^, is implicated in the metastasis of colon cancer Liu *et al*. ^64^ and was observed to be highly expressed in colorectal cancer cases Madan *et al*. ^25^. With mutated RNF43, as the Wnt signaling is enhanced, there is possibility of increased tumorigenesis and RRM2 could synergistically work in tandem with RAD51AP1 (another contributor of genomic stability) to enhance the metastasis of CRC. Coincidently, the pipeline gives a preferred low rank of 139 to RRM2-RAD51AP1 combination after the drug treatment, thus indicating that its inhibition in the suppressed cancer cells. The lower ranking of RRM2 along with the RNF43 points to the correct prioritization by the pipeline. Similar results were found for KI-67 (MKI67) which as an independent prognostic marker for CRC Melling *et al*. ^65^ and MCM4, which play essential role in DNA replication Nishihara *et al*. ^66^, are both highly expressed in colorectal cancer cases. AXIN2, like AXIN1, helps in the assembly of the destruction complex that facilitates in the degradation of *β*-catenin in the cytoplasm, thus negatively regulating the Wnt pathway Mazzoni and Fearon^67^. Mutations in AXIN family can lead to different subtypes of CRC. AXIN2 is also a transcriptional target for *β*-catenin and changes in protein levels of AXIN2 due to excessive *β*-catenin can negatively regulate the Wnt pathway in cancer cases. Mutations in RNF43 can lead to enhanced Wnt signaling which can then target the over expression of AXIN2 via *β*-catenin that might lead to negative feedback to the pathway at a later stage. After the ETC-1922159 treatment, AXIN2 was found to be down regulated. This implies that the drug is working at multiple levels to inhibit the Wnt pathway and the low ranking of the AXIN2-RNF43 combination assigned by the pipeline accounts for this fact.

ASCL2 has been found to play a major role in stemness in colon crypts and is implicated in colon cancer Zhu *et al*. ^68^. Switching off the ASCL2 leads to a literal blockage of the stemness process and vice versa. At the downstream level, ASCL2 is regulated by TCF4/*β*-catenin via non-coding RNA target named WiNTRLINC1 Giakountis *et al*. ^69^. Activation of ASCL2 leads to feedforward transcription of the non-coding RNA and thus a loop is formed which helps in the stemness and is highly effective in colon cancer. At the upstream level, ASCL2 is known act as a WNT/RSPONDIN switch that controls the stemness Schuijers *et al*. ^70^. It has been shown that removal of RSPO1 led to decrease in the Wnt signaling due to removal of the FZD receptors that led to reduced expression of ASCL2. Also, low levels of LGR5 were observed due to this phenomena. The opposite happened by increasing the RSPO1 levels. After the drug treatment, it was found that ASCL2 was highly suppressed pointing to the inhibition of stemness in the colorectal cancer cells. Also, Schuijers *et al*. ^70^ show that by genetically disrupting PORCN or inducing a PORCN inhibitor (like IWP-2), there is loss of stem cell markers like LGR5 and RNF43, which lead to disappearance of stem cells and moribund state of mice. A similar affect can be found with ETC-1922159, where there is suppression of RNF43 and LGR5 that lead to inhibition of the Wnt pathway and thus the ASCL2 regulation. These wet lab evidences are confirmed in the relatively low ranking of the combination ASCL2-RNF43 via the inhibition of PORCN-WNT that leads to blocking of the stemness that is induced by ASCL2. Since ASCL2 is directly mediated by the WNT proteins, the recorded ASCL2-WNT10B combination showed low priority ranking of 488, 497 and 321 for rbf, laplace and linear kernels, respectively, thus indicating a possible connection between WNT10B and ASCL2 activation. WNT10B might be playing a crucial role in stemness. This is further confirmed by wet lab experiments in Reddy *et al*. ^71^, which show BVES deletion results in amplified stem cell activity and Wnt signaling after radiation. WNT10B has been implicated in colorectal cancer Yoshikawa *et al*. ^72^.

The foregoing descriptions confirm the effectiveness of the pipeline for a few cases and it is not possible to elucidate each and every combination in a single manuscript. The rankings have been made available for further tests and will help the investigator in narrowing down their focus on particular aspects of the signaling pathway in cancer cases.

### Unidentified/(Un)identified 2^*nd*^-order combinations - one blade double edged sword

Hitherto, the 2^*nd*^ order combinations pertaining to known or recorded factors that have been affected by the ETC-1922159 have been prioritized and discussed. But there were some uncharacterized proteins that were also affected after the drug treatment and their behaviour might not be known in terms of higher order interactions. We now traverse the unchartered territory of unknown/unexplored/untested proteins whose transcript levels were recorded after the drug treatment. Note that we will be dealing with two different cases over here - (1) Unidentified/identified 2^*nd*^ order combinations and (2) Unidentified/unidentified 2^*nd*^ order combinations. In total 234 such unidentified proteins were recorded of which we show the interpretations of only a few. Note that the numbers like 1,13, 121 etc associated with XX DO NOT imply any sort of association and we do not assume anything about them. We just interpret the rankings of these unidentified factors with indentified and unidentified factors. Table 2 represents these combinations, with the first 20 describing XX1-identified factor combinations and the next 16 describing XX1-unidentified factor combinations.

**Table 2.** 2^*nd*^ order interaction ranking using HSIC for different kernels. Total number of interactions per gene were 2743.

| RANKING USING HSIC W.R.T XX1 |  |  |  |
| --- | --- | --- | --- |
| 2<sup>nd</sup> odr comb. | rbf | laplace | linear |
| UNIDENTIFIED-IDENTIFIED COMBINATIONS |  |  |  |
| XX1-RAD51AP1 | 199 | 434 | 36 |
| RRM2-XX1 | 131 | 185 | 138 |
| ASCL2-XX1 | 476 | 328 | 217 |
| XX1-AXIN2 | 552 | 331 | 419 |
| MKI67-XX1 | 67 | 299 | 48 |
| MCM3-XX1 | 188 | 3 | 26 |
| UBE2C-XX1 | 10 | 204 | 21 |
| XX1-LGR5 | 40 | 208 | 39 |
| XX1-PTPRO | 130 | 21 | 12 |
| XX1-NKD1 | 763 | 801 | 539 |
| XX1-HOXB8 | 740 | 178 | 411 |
| XX1-HOXB9 | 436 | 464 | 287 |
| XX1-PPA2 | 2399 | 2320 | 2659 |
| XX1-RNF43 | 2127 | 1488 | 2140 |
| XX1-XRCC6BP1 | 1475 | 1916 | 1732 |
| XX1-XRCC6 | 2569 | 2278 | 2734 |
| XX1-HOXB7 | 2533 | 2554 | 2708 |
| XX1-HOXB13 | 2631 | 2463 | 1579 |
| XX1-HOXA9 | 2006 | 1091 | 1823 |
| XX1-HOXA11 | 1860 | 1984 | 2326 |
| UNIDENTIFIED-UNIDENTIFIED COMBINATIONS |  |  |  |
| XX1-XX15 | 35 | 107 | 245 |
| XX1-XX110 | 112 | 763 | 101 |
| XX1-XX182 | 2734 | 2582 | 2669 |
| XX1-XX196 | 2689 | 2615 | 1964 |
| XX1-XX2 | 86 | 190 | 231 |
| XX1-XX20 | 105 | 386 | 15 |
| XX1-XX205 | 2669 | 2499 | 2610 |
| XX1-XX207 | 2671 | 2739 | 2733 |
| XX1-XX33 | 194 | 194 | 360 |
| XX1-XX35 | 430 | 600 | 654 |
| XX1-XX34 | 1969 | 1397 | 2173 |
| XX1-XX38 | 1968 | 1445 | 1509 |
| XX1-XX49 | 21 | 340 | 303 |
| XX1-XX47 | 115 | 814 | 119 |
| XX1-XX44 | 1690 | 465 | 1070 |
| XX1-XX45 | 1417 | 1830 | 2106 |

### unidentified-identified combinations

- We first begin with the analysis of the unknown factor XX1 with some of the identified and comfirmed factors that are implicated in the pathway either positively or negatively. As has been seen earlier many of the factors like RAD51AP1, RRM2, ASCL2, AXIN2, MKI67, LGR5 and NDK1 that are highly implicated in colorectal cancer case were found to be suppressed after the drug treatment. XX1 pairs with these factors and the pipeline assigns low rank to these combinations. One of the interpretations can be that XX1, like the above factors, is highly implicated in colorectal cancer case and its association with some of these factors might be possible in reality and entails verification of the same. For example, the minichromosome maintenance (MCM) proteins are essential replication factors and have been found to be overexpressed in colorectal cancer Ha *et al*. ^73^. After the administration of the drug, MCM3 was found to be highly suppressed and the pipeline points to its low rank along with XX1. Ubiquitin-conjugating enzyme E2C gene (UBE2C) Hao *et al*. ^74^ is known to be overexpressed in colorectal cancer Takahashi *et al*. ^75^ and its apparent downregulation after ETC-1922159 has been assigned a low rank with XX1. Genetic alterations in tyrosine phosphatases have been found in colorectal cancer and serve as tumor suppressors in colorectal cancer Laczmanska and Sasiadek^76^. PTPRO is one such family member and was found to be suppressed after drug treatment and assigned low ranks with XX1. There might be a possibility that mutations in PTPRO were present and there was overexpression of these mutated versions in the tumor cells before the drug administration. After the treatment, the in silico low ranks point to the observed suppressions in PTPRO. The HOX family, is known to play multiple roles in various tumor cases and varied affects have been found in colorectal tumor and normal cases Kanai *et al*. ^77^. HOXB8 and HOXB9 are highly expressed in colorectal cancer cases and were found to be downregulated after the drug treatment. Along with XX1, they were assigned low ranks as desired.

The opposite interpretations are that there are very high ranks assigned to many of the factors that are known to play tumor suppressor roles like HOXB7, HOXB13, HOXA9, HOXA11 Kanai *et al*. ^77^, PPA2 Cristóbal *et al*.^40^ and XRCC6BP1 and mutations in these could lead to enhancement in colorectal cancer. These were found to be down regulated after the drug treatment in cancer cases and were assigned high ranks along with the XX1. XX1 might be a tumor suppressor gene also that might be mutated in colorectal cancer case. But these high ranks provide an alternative insight to the functioning of XX1. Thus the interpretations are like two edges of one blade and biologists can use these rankings to see the multifaceted aspects of the hitherto unidentified XX1.

### unidentified-unidentified combinations

- Now we analyse the unknown factor XX1 with some of the unknown/unexplored factors that are implicated in the pathway either positively or negatively. Observing table 2, the rankings of unidentified-unidentified factors after the ETC-1922159 were also generated.

In each of the series starting with XXM, were *M* ∈ 1,2,3,4, two interactions are shown with very low ranks assigned to them and two interactions are shown with very high ranks assigned to them. For example, XX1-XX20 and XX1-XX2 both show low priority ranks while XX1-XX205 and XX1-XX207 high ranks. The low priority ranks indicate that their down regulation was important after the ETC-1922159 treatment and these might have been highly expressed in colorectal cancer case before the treatment of the drug. The high priority ranks indicate that these might be tumor suppressor genes (might be mutated) in colorectal cancer and would be highly expressed after the administration of the drug, had the mutations not happened in them. Their down regulation and yet high rank points to their mutated versions in the cancerous cells, a possibility that needs to be verified. Similar interpretations can be made for the different ranked unidentified-unidentified combinations that the pipeline generated on the list of downregulated genes after the ETC-1922159 treatment.

## Conclusion

Computational : A theoretically sound and a practical framework has been developed to prioritize higher order combinations of downregulated genes after the administration of ETC-1922159 PORCN-WNT inhibitor in cancer cells. The prioritization uses advanced density based sensitivity indices that exploit nonlinear relations in reproducing kernel hilbert spaces via kernel trick and support vector ranking method to rank and reveal various combinations of identified and unidentified factors that are affected after the drug treatment. This gives medical specialists/oncologists as well as biologists a way to navigate in a guided manner in a vast combinatorial search forest thus cutting down cost in time/investment/energy as well as avoid cherry picking unknown biological hypotheses.

Biological : MYC upregulates PRC2 complex. PRC2 complex contains EZH2, which suppresses tumor suppressor genes via epigenetic modifications. MYC & HOXB8 are up regulated in colorectal cancer, but dual working mechanism of the same is not known. The pipeline that is in silico showed ranking which correctly approximates and assigns low rank to this 3^*rd*^ order combination of MYC-HOXB8-EZH2, pointing to down regulation by ETC-1922159. If the protein interaction and MYC-HOXB8 can be established and a study be done apropos EZH2, it will establish at in vitro/in vivo level, the in silico ranking also.

## Conflict of interest

There are no conflicts to declare.

## Dedication

The author dedicates this small work to his long time friend Joana Carolina Martins who was diagnosed with tumor at the age of 35 and to other cancer patients. Joana's diagnosis as well as communications propelled the author to delve into the research work related to cancer biology and therapeutics.

## Author’s contributions

SS designed, developed and implemented the insilico experimental setup, wrote the code, generated and analysed the results and wrote the manuscript.

## Acknowledgements

Special thanks to Mrs. Rita Sinha and Mr. Prabhat Sinha for supporting the author financially, without which this work could not have been made possible. Marco Wiering and Silja Renooij for continued support during the years of independent research work.

## Code Availability

Code on google drive at https://drive.google.com/drive/folders/0B7Kkv8wlhPU-WDgzdUVfTzA2cW8 Some of the 2^*nd*^ order rankings in https://drive.google.com/file/d/0B7Kkv8wlhPU-UkVNVkx3TzNQcU0/view

## Source of Data

Data used in this research work was released in a publication in Madan *et al*. ^25^. The ETC-1922159 was released in Singapore in July 2015 under the flagship of the Agency for Science, Technology and Research (A*STAR) and Duke-National University of Singapore Graduate Medical School (Duke-NUS).

## Appendix

### Choice of sensitivity indices

The SENSITIVITY PACKAGE (Faivre *et al*.^29^ and Iooss and Lemaître^30^) in R langauge provides a range of functions to compute the indices and the following indices will be taken into account for addressing the posed questions in this manuscript.

1. sensiFdiv - conducts a density-based sensitivity analysis where the impact of an input variable is defined in terms of dissimilarity between the original output density function and the output density function when the input variable is fixed. The dissimilarity between density functions is measured with Csiszar f-divergences. Estimation is performed through kernel density estimation and the function kde of the package ks. Borgonovo^78^ and Da Veiga^79^
2. sensiHSIC - conducts a sensitivity analysis where the impact of an input variable is defined in terms of the distance between the input/output joint probability distribution and the product of their marginals when they are embedded in a Reproducing Kernel Hilbert Space (RKHS). This distance corresponds to HSIC proposed by Gretton *et al*. ^80^ and serves as a dependence measure between random variables.
3. soboljansen - implements the Monte Carlo estimation of the Sobol indices for both first-order and total indices at the same time (all together 2p indices), at a total cost of (p+2) × n model evaluations. These are called the Jansen estimators. Jansen^81^ and Saltelli *et al*.^82^
4. sobol2002 - implements the Monte Carlo estimation of the Sobol indices for both first-order and total indices at the same time (all together 2p indices), at a total cost of (p+2) ×n model evaluations. These are called the Saltelli estimators. This estimator suffers from a conditioning problem when estimating the variances behind the indices computations. This can seriously affect the Sobol indices estimates in case of largely non-centered output. To avoid this effect, you have to center the model output before applying “sobol2002”. Functions “soboljansen” and “sobolmartinez” do not suffer from this problem. Saltelli^83^
5. sobol2007 - implements the Monte Carlo estimation of the Sobol indices for both first-order and total indices at the same time (all together 2p indices), at a total cost of (p+2) × n model evaluations. These are called the Mauntz estimators. Saltelli and Annoni^84^
6. sobolmartinez - implements the Monte Carlo estimation of the Sobol indices for both first-order and total indices using correlation coefficients-based formulas, at a total cost of (p + 2) × n model evaluations. These are called the Martinez estimators.
7. sobol - implements the Monte Carlo estimation of the Sobol sensitivity indices. Allows the estimation of the indices of the variance decomposition up to a given order, at a total cost of (N + 1) × n where N is the number of indices to estimate. Sobol’^85^

